# Kinetic structure of recycling vesicle pool at the Calyx of Held synapse under in vivo-like activities

**DOI:** 10.1101/2021.03.03.433702

**Authors:** Zili Liu, Ying Zhu, Yubing Hu, Jianyuan Sun

## Abstract

Synaptic transmission at mammalian central synapses has ongoing background activity at physiological temperature. The recycling vesicle pool, with proper kinetics, ensures sustained synaptic transmission. However, the kinetic structure of recycling vesicle pool has never been quantitatively analyzed before, and most studies were performed at room temperature and under resting conditions. With the combination of presynaptic capacitance measurement and postsynaptic EPSC recording on calyx of Held synapses at physiological temperature, we studied vesicle recycling under sustained presynaptic stimulation. The kinetics of vesicle reuse was revealed by impeding transmitter refilling with folimycin. We kinetically dissected the recycling vesicle pool as sequentially connected sub-pools and depicted the complete kinetic structure. The sizes and transition rates among these sub-pools were dynamically regulated by neuronal activity, in order to ensure efficient synaptic transmission. Our work highlights the impact of the vesicle recycling machinery on stable and reliable synaptic transmission under variable levels of neuronal activity.

**Impact statement:** The recycling pool of vesicles are kinetically dissected as four populated pools ensuring stable and reliable synaptic transmission

## Introduction

Synaptic vesicles (SVs) recycle locally within the presynaptic terminal for repeated release (Denker & Rizzoli, 2010; Rizzoli & Betz, 2005; Soykan, Maritzen, & Haucke, 2016; Südhof, 2004). Conceptually, SVs undergo recycling through the stages of priming, exocytosis, endocytosis and intra-terminal trafficking including cargo sorting, and rapid refilling of newly formed SVs with neurotransmitter. To maintain synaptic efficacy, an efficient kinetic structure is required to support persistent consumption and reconstitution of SVs (Alabi & Tsien, 2012; Mohrmann, De Wit, Verhage, Neher, & Sørensen, 2010; Qiu, Zhu, & Sun, 2015). Several mechanisms of SV recycling mediated via clathrin mediated endocytosis (CME), and clathrin independent ultrafast endocytosis, kiss-and-run and activity-dependent bulk endocytosis (ADBE) were proposed depending on activity levels or synapse types (Smith, Renden, & von Gersdorff, 2008; Watanabe & Boucrot, 2017; Watanabe et al., 2014; L. G. Wu, 2004). However, categorization of these modes of SV recycling has mostly been based on SV endocytosis, with little consideration about post-endocytic SV trafficking. Hundreds to many thousands of SVs exist in a presynaptic terminal and a large proportion of them participate in recycling, and are defined as the recycling pool (RP) (Denker & Rizzoli, 2010; Qiu et al., 2015; Rizzoli & Betz, 2005; Südhof, 2004). Significant heterogeneity in spatial distribution and competence of reuse was revealed among SVs in RP (Denker & Rizzoli, 2010; Qiu et al., 2015). Apart from the readily releasable pool (RRP) that has been well defined and assayed, it is still unclear how the residing SVs in nerve terminals are kinetically organized for synaptic transmission. It was reported that SV proteins such as VAMP/synaptobrevin and synaptotagmin were preferentially recruited from the reservoir at the surface of the nerve terminal, but it remains unclear whether a corresponding surface pool of SV membrane is readily retrievable at the putative sites of endocytosis (Fernández-Alfonso, Kwan, & Ryan, 2006; Hua et al., 2011; Wienisch & Klingauf, 2006). Recent studies show that the SVs in a nerve terminal are kinetically composed of multiple pools with different timing and preference of reuse (Miki, Nakamura, Malagon, Neher, & Marty, 2018; Qiu et al., 2015). By quantitatively analyzing the kinetics of RP depletion from resting status, two sequentially connected sub-populations of SVs before priming into RRP were defined (Qiu et al., 2015). Importantly, all of these relevant observations were made from a resting status and at room temperature (RT). Although temperature has been found to influence SV exocytosis and endocytosis, it still remains to be elucidated whether temperature functionally affects the intra-terminal SV trafficking (Hoopmann et al., 2010; Renden & von Gersdorff, 2007; Sabatini & Regehr, 1996; Watanabe et al., 2014). It was found that ultrafast endocytosis was predominant under *in vivo*-like stimulation, corresponding to both brief neuronal activity and high-frequency stimulation at physiological temperature (PT) (Soykan et al., 2017; Watanabe et al., 2014). More impressively, the ultrafast endocytosis generates the endocytic SVs which have larger size than exocytotic SVs without the requirement for clathrin, implying temperature can regulate SV recycling in not only kinetics, but also molecular perspectives (Chanaday & Kavalali, 2018; Delvendahl, Vyleta, von Gersdorff, & Hallermann, 2016; Kusick et al., 2018; Watanabe & Boucrot, 2017; Watanabe et al., 2014). Furthermore, the neuronal activities *in vivo* are highly dynamic, and synaptic signals are often embedded in a wide range of stochastic background (Geisler, Deng, & Greenberg, 1985; Hermann, Pecka, von Gersdorff, Grothe, & Klug, 2007; Liberman, 1978; Sommer, Lingenhöhl, & Friauf, 1993), which gives rise to a pivotal question of whether and how SVs recycling machinery shapes synaptic signaling and short-term plasticity in addition to supplying release-competent SVs? Here, we addressed these questions by dissecting the kinetic structure of SV recycling. With the combination of presynaptic capacitance measurement and evoked excitatory postsynaptic current (EPSCs) recordings on the calyx of Held synapse under *in vivo*-like activities, we quantitatively analyzed vesicle recycling. We found that the whole RP kinetics could be divided into four populated pools which were sequentially connected and functionally acted as regulatory stages along the recycling pathway. The combined kinetic structure of the RP could function as not only an efficient SV provider, but also a stabilizer of synaptic transmission.

## Results

### Steady state of synaptic vesicle release and RRP during sustained stimulations

Different from most of the previous studies in resting status at RT, we studied synaptic vesicle recycling under *in vivo*-like activities, i.e. introducing 20 Hz or higher frequency presynaptic firing to *in vitro* acute mouse brain slice at near physiological temperature (Hermann et al., 2007; Renden & von Gersdorff, 2007; Watanabe et al., 2014). We found that the miniature excitatory postsynaptic currents (mEPSCs) at PT had larger amplitude, faster kinetics and higher frequency in comparison to mEPSCs at RT (Figures S1A–S1F), displaying a significant temperature-dependence in spontaneous vesicle release.

We recorded the postsynaptic response to 20 Hz temporally Poisson-distributed afferent fiber stimulation which mimicked the physiological background firing at the calyx of Held synapse *in vivo* (Hermann et al., 2007). This mimicking stimulation (240 s) along with the following uniform stimulation (120 s) at fundamental frequency of 20 Hz was applied to the recorded synapses (Figures S2A and S2C). No significant difference in EPSC amplitudes at steady state was found corresponding to these stimulations with little change in the variation of amplitudes, indicating the similar vesicle supply under both stimulations (Figures S2B, S2D and S2E). Therefore, the uniform stimulation was applied in the rest of our study.

In our study, 50 Hz stimulation was applied to mimic the response to acoustic stimulation induced afferent input to this synapse (Sommer et al., 1993). In comparison to 20 Hz stimulation, EPSCs induced by 50 Hz stimulation showed more significant depression with shorter time to reach steady state at physiological temperature (Figures 1A–1E) and room temperature (Figures S1G–S1K). The EPSCs recorded at PT displayed larger absolute amplitudes and quantal contents (as EPSC/mEPSC) and longer time to reach the steady state than EPSCs recorded at RT, indicating the significantly increased transmitter consuming which required the enhanced vesicle supply during neural activities at PT (Figures 1F and 1G).

**Figure 1.**
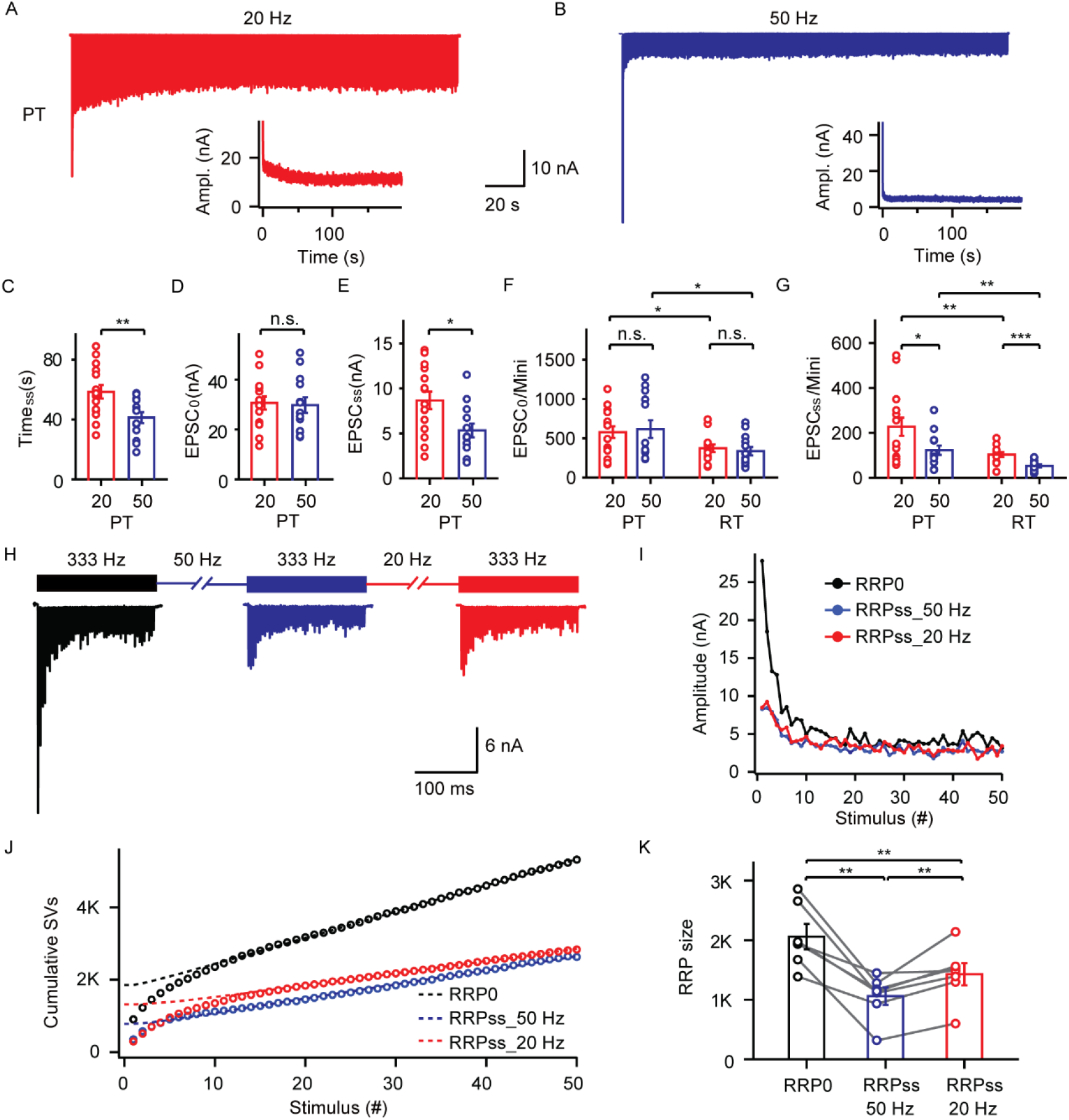
Steady state of synaptic transmission and RRP measurement under sustained stimulations. (A and B) Representative EPSCs and the corresponding amplitudes during sustained stimulation at 20 Hz (A) or 50 Hz (B) at physiological temperature. (C-E) Statistics of steady state time (C), first EPSC amplitude (D), and steady state amplitude (E) of each train at 20 Hz or 50 Hz stimulation at physiological temperature (20 Hz: n=15; 50 Hz: n=13). (F and G) Statistics of the quantal contents (EPSC/mEPSC) of first EPSC (F) and steady state EPSC (G) between 20 Hz and 50 Hz at RT or PT. (H) Protocol for estimation of RRP at resting status and steady state under 20/50 Hz fiber stimulation and the representative records. (I) Representative plot of EPSCs amplitude versus stimulus number at 333 Hz at resting status and steady state under 20/50 Hz fiber stimulation corresponding to (H). (J) Representative plot of cumulative vesicle number versus stimulus number at 333 Hz and instant RRP size at resting status and steady state under 20/50 Hz fiber stimulation. (K) Statistics of RRP size at resting status, steady state (RRP_ss_) under 20 Hz/50 Hz stimulation, respectively (n = 7). Data are represented as mean ± SEM. \**p* < 0.05, \*\**p* < 0.01, \*\*\**p* < 0.001.

To determine the immediately available vesicle supply under different stimulations, we evaluated the instant RRP. In the same synapse, a train of 50 stimuli at 333 Hz was applied to measure the RRP instantly at resting status and the RRP during steady state conditions (RRP_ss_) for 20 Hz or 50 Hz stimulation (Figures 1H–1J) (Thanawala & Regehr, 2013; L. G. Wu & Borst, 1999). The RRP_ss_ size for 20 Hz stimulation was estimated as 1427 ±186 (n = 7) while the RRP_ss_ for 50 Hz stimulation contained 1059 ± 148 vesicles. These steady state RRPs were significantly smaller than the RRP at resting status (2059 ± 213) (Figure 1K). Thus, RRP size was dynamically regulated by the level of neuronal activity.

### Kinetics of vesicle reuse revealed by impeding transmitter refilling with folimycin

To investigate the kinetics of vesicle recycling, we locally applied folimycin (4 μM), a vacuolar-ATPase inhibitor, to impede vesicle reacidification and transmitter refilling when the EPSCs amplitude reached steady state under sustained stimulation (Figure 2A) (Qiu et al., 2015). After folimycin perfusion, the amplitude of the EPSCs was significantly decreased but not eliminated (Figure 2B). We attribute the suppression of EPSCs to the inhibition of neurotransmitter refilling by folimycin and the unexhausted EPSCs to the insufficiency of pharmacological application of folimycin. EPSCs were expected to reach a new steady state with much-reduced amplitude when all of the preserved vesicles (not reused yet) were depleted and the newly endocytic vesicles were fully reused (Qiu et al., 2015). Because several-minute application of folimycin does not affect RRP and release probability assayed by capacitance measurement, the decrease of EPSC amplitudes has to be proportional to the reduction of mEPSC amplitudes (Qiu et al., 2015). The quantal size of these reused vesicles (q_foli_) was thus estimated as 4.69 pA, ~10 fold smaller than normal one (q_ctrl_ = 48.92 pA, n = 8) (Figure 2C; See Eq.1 in STAR Methods). We took the cumulative amplitude of the recorded EPSCs as a measure of the released transmitter from exocytosed vesicles (Figure 2D and 2F). During folimycin application, the nerve terminal was consuming two populations of vesicles, the preserved vesicles which were alreadyexisting in the terminal and not affected by folimycin (preserved SVs; the summed number was N_preserved_) and the reused vesicles (reused SVs; the summed number was N_reused_) (Qiu et al., 2015). Meanwhile, the amplitude of EPSCs at the steady state were expected to keep constant if folimycin was not applied (Figure 2B dash line) and the cumulative EPSC amplitude was attributed to the sum of the released SVs (N_preserved_ and N_reused_) with q_ctrl_ (see Eq.2 in STAR Methods). Comparatively, both N_preserved_ with q_ctrl_ and N_reused_ with q_foli_ contributed to the cumulative EPSC amplitude in the presence of folimycin (See Eq.3 in STAR Methods). As derived in Eq.2 and Eq.3, N_preserved_ and N_reused_ at any time during stimulation could be estimated and we thus completed the analysis of the kinetics of vesicle recycling (Figure 2E and 2G) (Qiu et al., 2015). It was estimated that the cumulative number of released preserved SVs under sustained 20 Hz or 50 Hz stimulation tended to be unchanged at the end of prolonged stimulation, indicating the depletion of preserved SVs. The total preserved vesicles number could be counted as the ultimate value of the cumulative curve, 263,213 ±13,112 (n = 7) for 20 Hz stimulation and 259,866 ± 21,897 (n = 8) for 50 Hz stimulation respectively, with no significant difference (*p* > *0.9*) (Figure 2E and 2G). Correspondingly, the fastest reuse time for the newly endocytosed vesicles was estimated as 10.97 ± 0.84 s when 8.91% ± 1.65% of the preserved vesicles are released during 20 Hz stimulation, and 6.95 ±1.07 s when 7.89% ± 1.20% of the preserved vesicles are released during 50 Hz stimulation (Figure S3). Thus, we estimated the intra-terminal vesicle numbers and minimum reuse time under different frequencies of sustained stimulation at PT by analyzing the kinetics of vesicle recycling.

**Figure 2.**
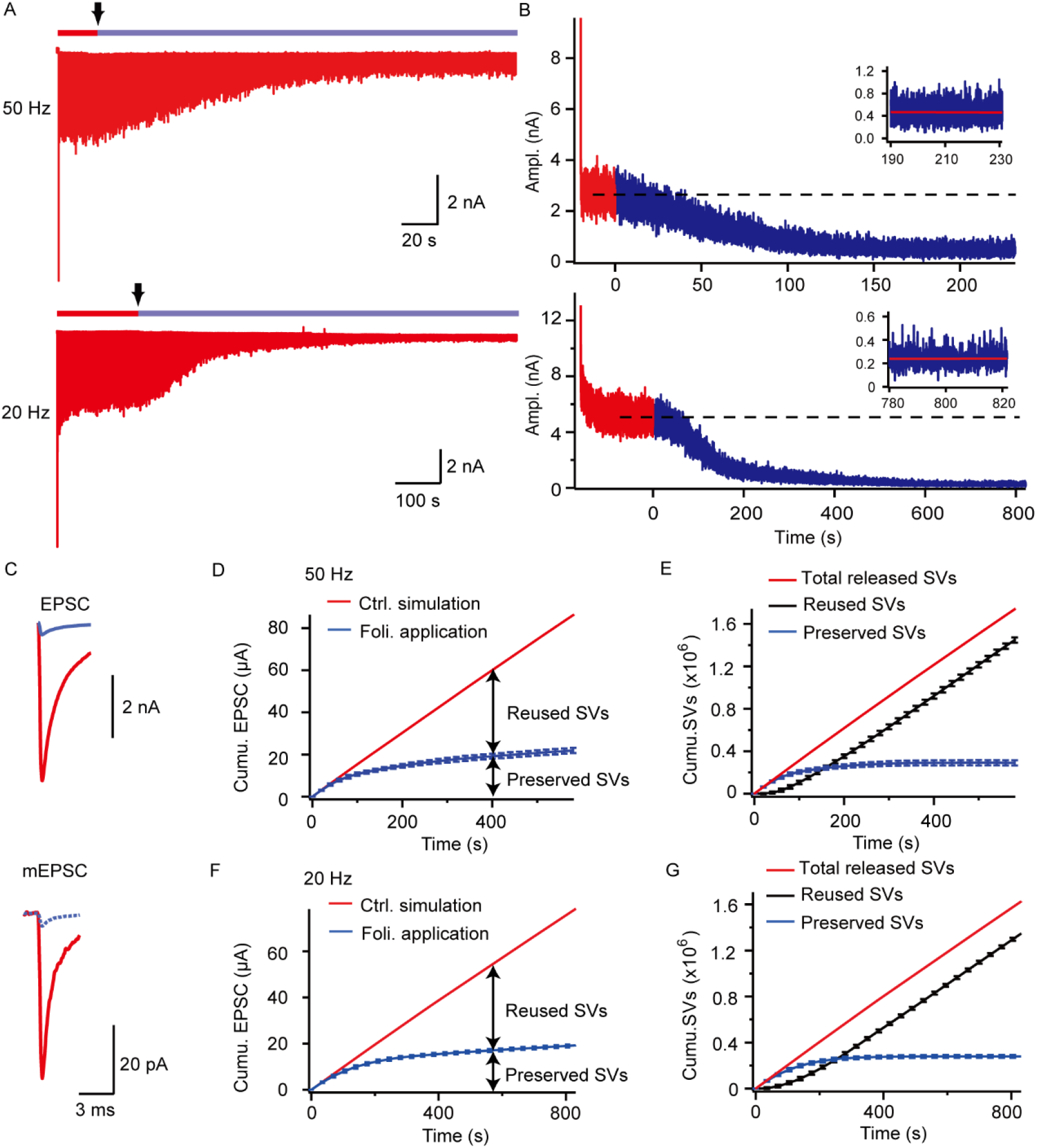
Kinetics of vesicle reuse revealed by impeding transmitter refilling with folimycin. (A) Representative EPSCs corresponding to 50 Hz (Upper) or 20 Hz (Lower) stimulation before and after folimycin application and the time point of folimycin application was indicated by the black arrow on the top of experimental strategy bar. (B) The EPSC amplitudes corresponding to (A) without (red) and with (blue) folimycin application; the averaged amplitude of EPSCs at the steady state was demonstrated by black dish line. (Inset) EPSC amplitudes in the last 40 s of stimulation with folimycin fitted with linear regression (red line). (C) The steady state EPSC before and after folimycin application (Upper); the recorded mEPSC before folimycin application and estimated mEPSC after folimycin application (Lower). (D and F) Cumulative EPSC amplitudes under 50 Hz stimulation (n = 8) (D) and 20 Hz stimulation (n = 7) (F) with (blue) and without folimycin (red). (E and G) Calculated cumulative number of total released vesicles (red), preserved vesicles (blue), and reused vesicles (black) under 50 Hz stimulation (n = 8) (E) and 20 Hz stimulation (n = 7) (G).

### The kinetic structure of synaptic vesicles in calyceal terminal during sustained stimulations

We differentiated the preserved vesicle depletion curve and got the depletion rate (R_depl_). Similar to the result obtained at RT, R_depl_ could be fitted by a double exponential function (Figure 3A and 3B, See Eq.11 in STAR Methods) (Qiu et al., 2015). Accordingly, we proposed that the recycling pool was composed of the RRP, the readily priming pool (RPP), and the following pool (FP) with k_1_ and k_2_ as the corresponding supply rate (Figures 3C and 3D) (Qiu et al., 2015). The kinetic process could be described as a group of differential equations with the assumption that during steady state conditions, the SV release rate was the same as the priming rate. Before folimycin application, RRP_ss_, RPP and FP were all occupied by the preserved vesicles. The three-pool model well described the RP depletion kinetics (Figures 3A and 3B, white curve), and the size of RPP and the FP at steady state were thus estimated as 119,709 ± 12,342 and 142,339 ± 9,877 at 20 Hz; 69,748 ± 12,337 and 189,238 ± 32,312 at 50 Hz by fitting R_depl_ with the analytic solution of these equations (See Eq.11 and Eq.12 in STAR Methods).

**Figure 3.**
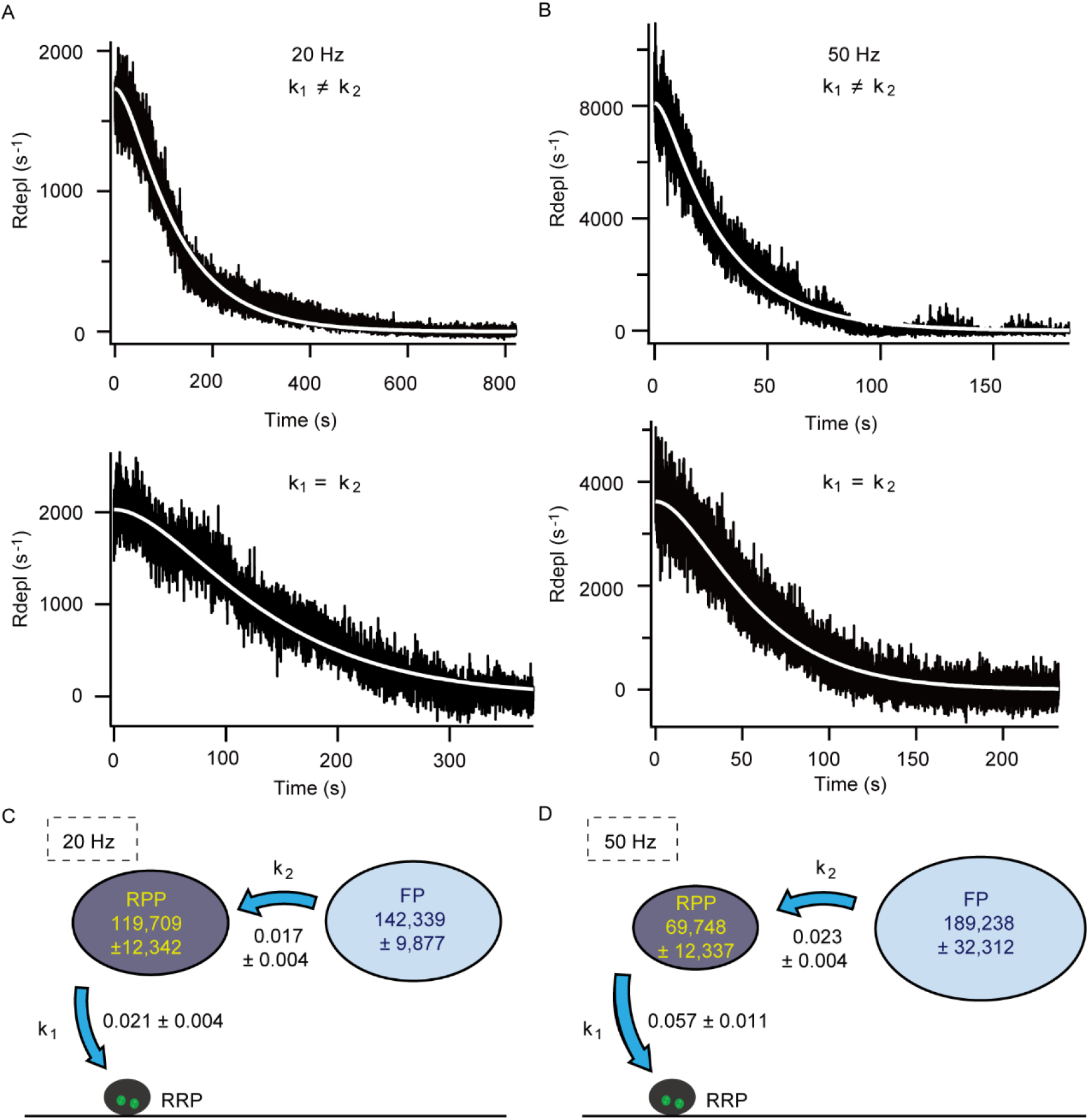
The kinetics of preserved synaptic vesicle depletion at the calyceal terminal during sustained stimulations. (A and B) Representative preserved pool depletion rate (R_depl_) trace under 20 Hz stimulation (A) and 50 Hz stimulation (B) fitted with double-exponential fitting (k_1_ ≠ k_2_, Upper) or mono-exponential fitting (k_1_ = k_2_ ≈ k, Lower). (C and D) Quantification of dissected synaptic vesicle pools and transition rates by three pool depletion model under 20 Hz stimulation (n = 7) (C) and 50 Hz stimulation (n = 8) (D).

To validate that a three-pool model is sufficient, we used a hypothetic three-plus-pool model, assigning two pools (FP1, FP2) following RPP, to describe the kinetics of preserved vesicle depletion (Figures S4A-D). Corresponding differential equations were derived to fit R_depl_ and we estimated the sizes of each sub-pool analytically. Interestingly, the size of FP_2_ (4,575 ± 1,039 at 20 Hz and 2,100 ± 461 at 50 Hz) was relatively small compared with FP_1_ and the transition rate from FP_2_ to FP_1_ (0.866 ± 0.377 at 20 Hz and 2.086 ± 0.366 at 50 Hz) was much larger than the transition rate from FP_1_ to RPP (0.017 ± 0.004 at 20 Hz and 0.022 ± 0.004 at 50 Hz), strongly suggesting that FP_2_ to FP_1_ could be kinetically merged as a single pool (Figures S4C and S4D). This notion was consistently supported by the similar SD of three-pool model and three-plus-pool model fitting (SD = 325.32 ± 62.93 and 323.69 ± 62.80, respectively). As FP was the last stage for the preserved pool depletion, this pool was considered as the first kinetic population for gathering endocytic vesicles and we will now refer to this as the post-endocytic pool (PEP).

By quantitatively analyzing the depletion of the preserved pool at steady states under sustained stimulation, we revealed that intra-terminal vesicles can be kinetically depicted as a sequential three pool system.

### Kinetic structure of recycling vesicle pool under sustained stimulations

A significant amount of the vesicular molecules are found stranded at the surface of nerve terminals during neuronal activity, suggesting the existence of a surface pool of vesicles (Fernández-Alfonso et al., 2006; Hua et al., 2011; Wienisch & Klingauf, 2006). To determine whether the putative vesicular membranes are accumulated at the surface of nerve terminal, we made capacitance measurements of the calyceal terminal in response to trains of action potential-like stimuli (1ms depolarization from −80 mV to + 40 mV) at a frequency of 20 Hz or 50 Hz (Figure 4A). The train of stimulation caused a dramatic increase of membrane capacitance which tended to reach a steady state, indicating a population of SV membrane accumulated at the terminal and formed a surface pool (SP). Under 50 Hz stimulation, the SP size, as membrane capacitance change, was 2.38 ± 0.38 pF (n = 9), which was significantly larger than the SP size under 20 Hz stimulation (1.75 ± 0.15 pF, n = 9, *p* = 0.009) (Figures 4B and 4C). As the average capacitance of a single SV membrane is ~65 aF, 36,600 and 26,900 vesicles were estimated in the SP under sustained 50 Hz and 20 Hz stimulation, respectively (Figure 4D) (J. Y. Sun, Wu, & Wu, 2002).

**Figure 4.**
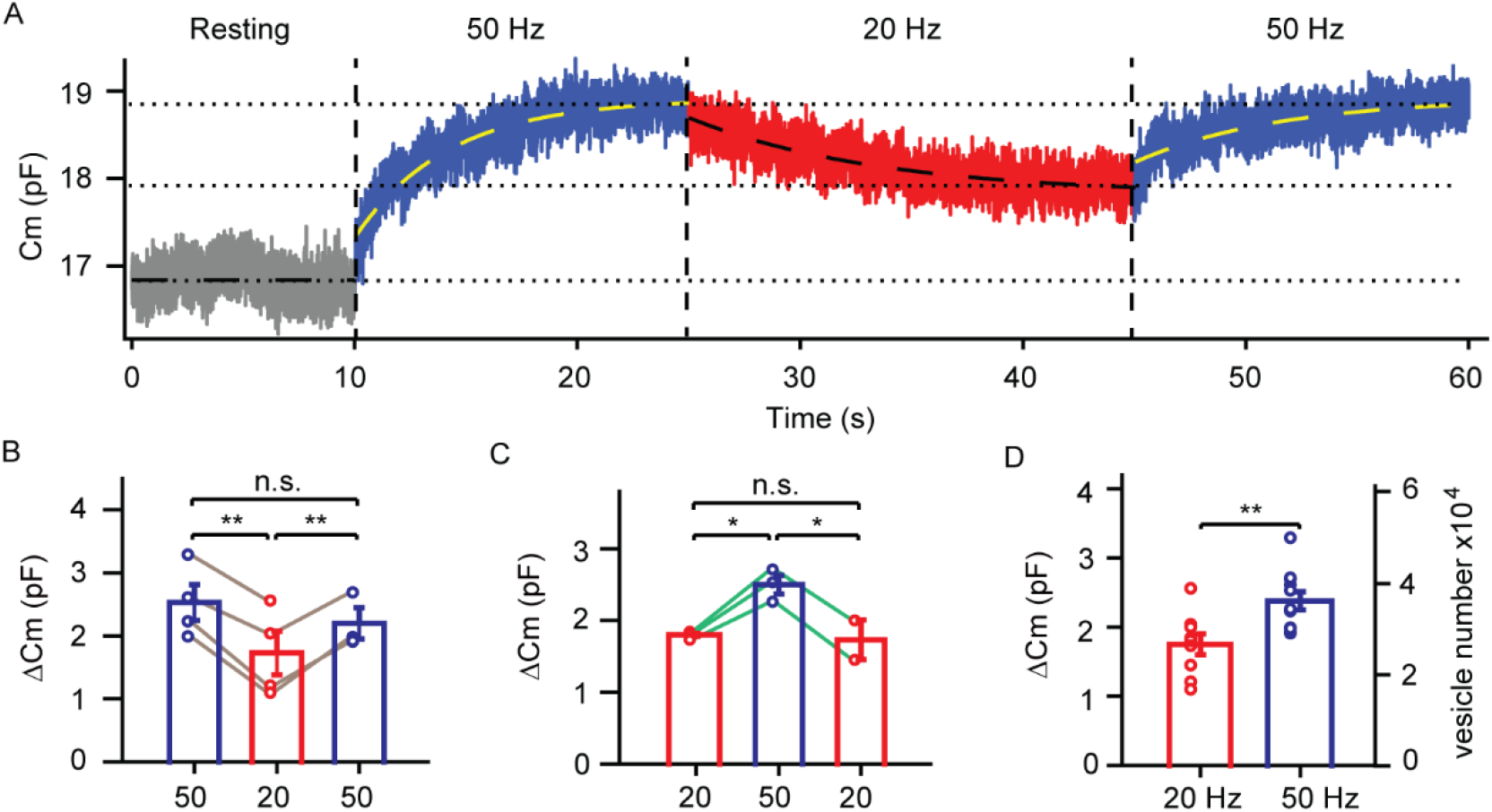
The surface pool of synaptic vesicles under sustained stimulations. (A) Representative surface pool measurement during 20 and 50 Hz presynaptic AP-like stimulation; the tansient process was fitted with mono-exponental function. (B) Statistics of the capacitance change induced by the continuous depolarization of 50 Hz-20 Hz− 50 Hz sequence. (C) Statistics of the capacitance change induced by the continuous depolarization of 20 Hz-50 Hz- 20 Hz sequence. (D) Quantification of the capacitance change and the corresponding surface pool size under 20 and 50 Hz presynaptic stimulation (20 Hz: n = 9; 50 Hz: n = 11). Data are represented as mean ± SEM. \**p* < 0.05, \*\**p* < 0.01.

Given the SP together with the intra-terminal three pools, a four-pool model was thus applied to describe the kinetics of vesicle recycling under sustained neuronal activity. The steady state of SP under a prolonged stimulation resulted from the equilibrium of exocytosis and endocytosis (Figure 4A; Figure S6A). We thus estimated the vesicle release rate (k_0_) as the stimulation frequency × EPSC/RRP_ss_ and the vesicle retrieval rate (k_3_) as frequency × EPSC/SP. By normalization, the RRP_ss_, RPP, PEP and SP occupied 0.49 ± 0.06%, 41.22 ± 4.25%, 49.01 ± 3.40% and 9.28 ± 0.79% of the overall RP vesicles under 20 Hz stimulation; and 0.36 ± 0.05%, 23.52 ±4.16%, 63.80 ± 10.89% and 12.32 ± 0.68% correspondingly under 50 Hz stimulation (Figure 5). The inter-pool transition rates were also estimated, as k_0_ = 3.216 ± 0.584, k_1_ = 0.021 ± 0.004, k_2_ = 0.017 ± 0.004, k_3_ = 0.170 ± 0.031 under 20 Hz stimulation and k_0_ = 9.654 ± 1.190, k_1_ = 0.057 ± 0.011, k_2_ = 0.023 ± 0.004, k_3_ = 0.280 ± 0.034 under 50 Hz stimulation (Figure 5). Therefore, we completely dissected the kinetic structure of RP under *in vivo*-like neuronal activity levels.

**Figure 5.**
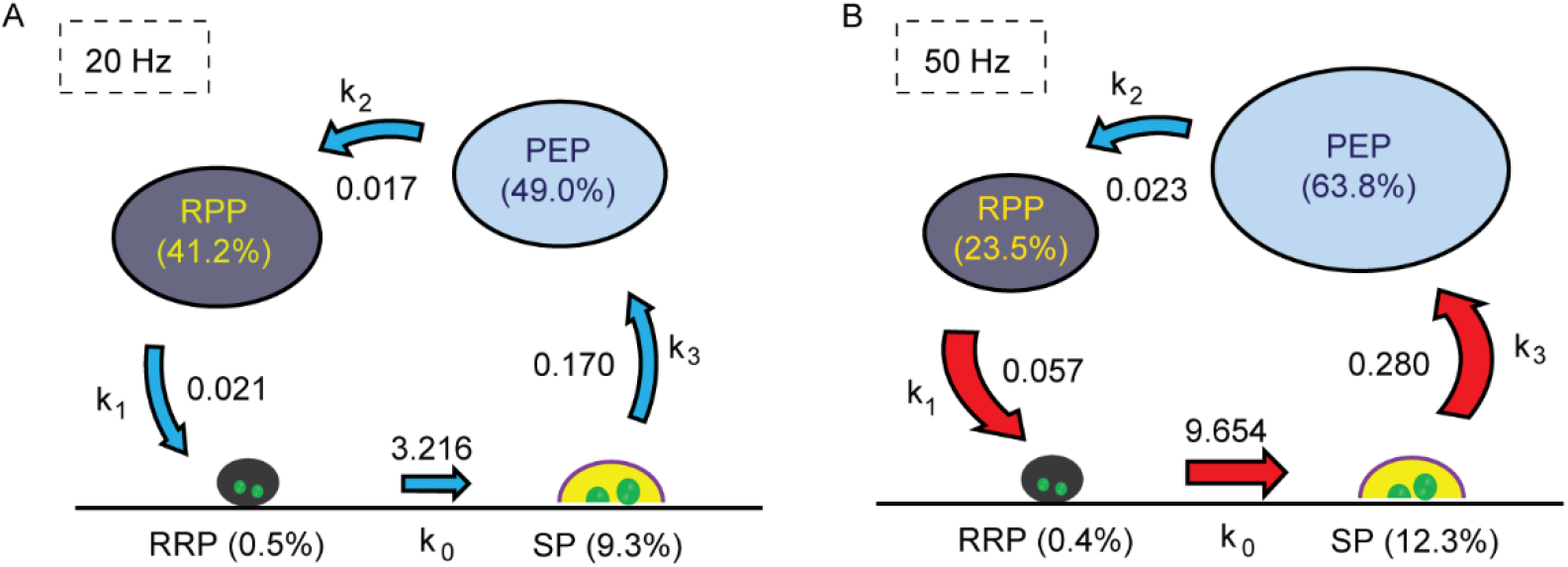
Kinetic structure of recycling vesicle pool under sustained stimulations. (A and B) Quantification of dissected synaptic vesicle pools and inter-pool transition rates in terms of total recycling vesicles in the calyceal terminal at 20 Hz (n = 7) (A) and 50 Hz (n = 8) (B) stimulation.

### Synaptic transmission under variant neuronal activities

In the MNTB, the presynaptic firing frequency varies over a range of hundreds of hertz periodically embedded in the background firing at ~20 Hz corresponding to various acoustic input (Hermann et al., 2007; Sommer et al., 1993). The kinetic structure of vesicle recycling is required to ensure not only the vesicle supply but also the robustness of synaptic signaling under sustained and variant neuronal activities. To study this, we designed an experiment to test whether/how much the calyx of Held synapse could maintain a similar synaptic response to the same synaptic inputs embedded background firing. We recorded the EPSCs in response to the stimulation with frequency-switchover between 20 Hz and 50 Hz at different durations. We first gave a long train of 20 Hz stimulation as background firing and recorded the EPSC responses. When the postsynaptic response reached a steady state, the stimulation frequency was then switched to 50 Hz for 5 s, 30 s and 90 s and back (Figures 6A–6C). In order to directly compare the postsynaptic responses corresponding to 20 Hz and 50 Hz stimulation, the release rate was calculated as the cumulative EPSC amplitudes per 100 ms and then normalized. We found a brief switchover (duration < 5 s) to 50 Hz did not significantly change the synaptic response to 20 Hz stimulation (Figures 6D, 6G–6H). When the duration of frequency switchover was prolonged (10 s or longer), a decrease in the release rate upon at 20 Hz stimulation was observed (Figures 6D–6H). This deviation diminished after certain period of time depending on the duration of frequency switchover (Fig. 6D-H). We attributed these phenomena to the intrinsic property of recycling kinetics as our kinetic model could well predict all of the observations (Black curves in Figures 6D–6G; Red curve in Figure 6H). Both the experiment and modelling indicate that the kinetic structure of vesicle recycling was able to maintain the stability of the vesicle supply under brief (< 5 s) stimulation frequency fluctuations (20 Hz − 50 Hz).

**Figure 6.**
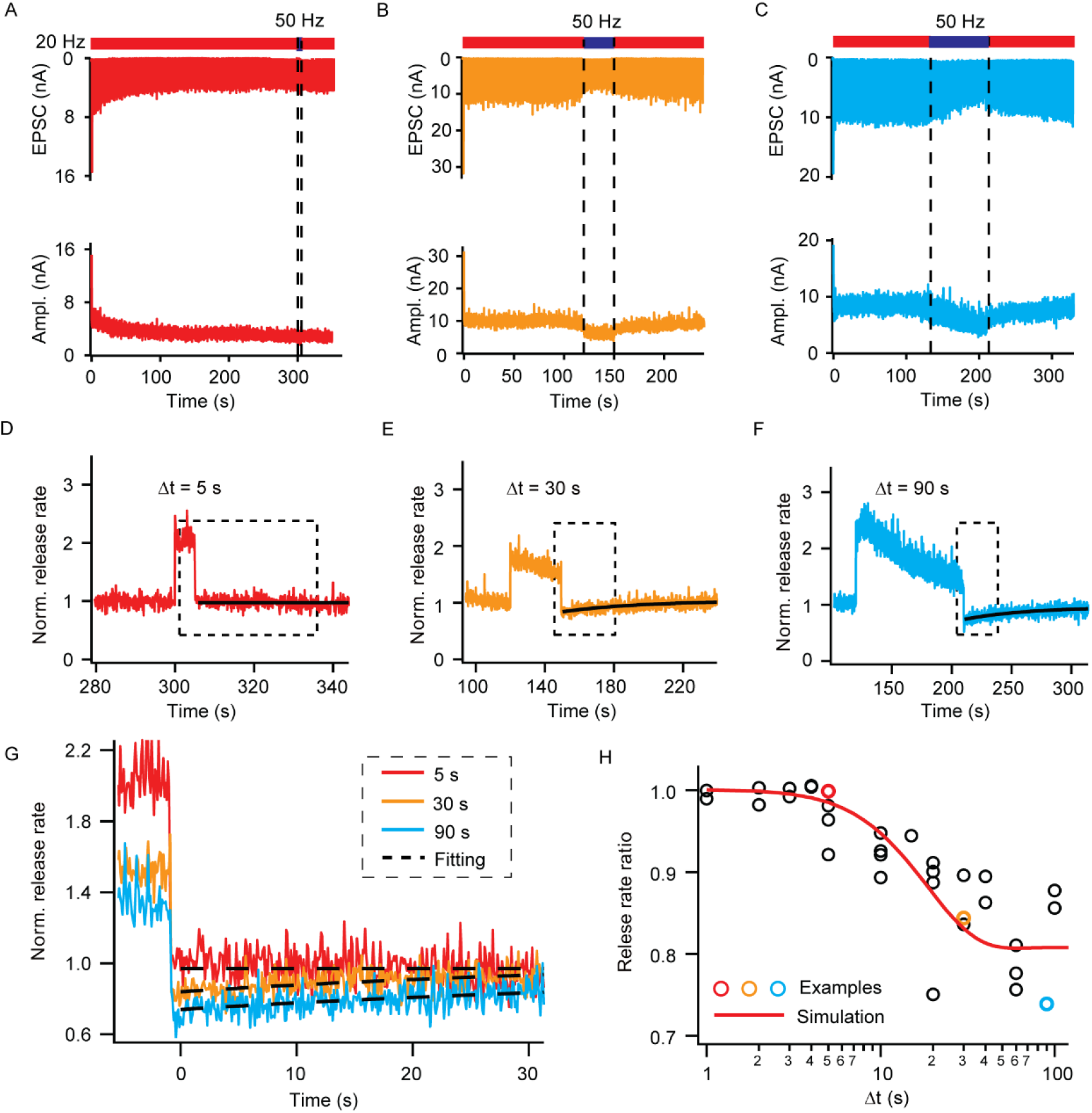
Synaptic transmission under variant neuronal activities. (A-C) Representative recorded EPSCs and the corresponding amplitudes in response to the stimulation with frequency-switchover between 20 Hz and 50 Hz in 5 s (A), 30 s (B) and 90 s (C) after the postsynaptic response reached a ready state at 20 Hz stimulation. (D-F) The corresponding normalized cumulated EPSC amplitudes per 100 ms during the frequency-switchover in A-C. (G) Local zoom of the representative normalized cumulated EPSC amplitudes in dashed box corresponding to D-F (H) Plot of the relative release rate versus frequency-switchover interval at 20 Hz stimulation and the best fit result.

## Discussion

### Synaptic vesicle exocytosis and endocytosis under *in vivo*-like neural activities

We observed the kinetics of vesicle recycling under *in vivo*–like conditions. All the observations were made under physiological temperature. Consistent with previous reports, mEPSCs had higher frequency and larger amplitude with accelerated kinetics, and evoked EPSCs displayed larger size in comparison to the recording at RT (Figures 1C–1E; Figures S1A–S1K) (Kuba, Yamada, & Ohmori, 2003; Kushmerick, 2006; Micheva & Smith, 2005; Sabatini & Regehr, 1996; Samigullin, Bukharaeva, Nikolsky, & Vyskočil, 2003). Notably, the ratio of electrical charge of EPSC and mEPSC at PT was significantly increased, indicating single presynaptic impulse induced more vesicle release, which required more vesicle supply at PT (Figures 1F and 1G). As the RRP sizes were similar at PT and RT, we determined that the enhanced vesicle release at PT was due to an increase in the release probability (Qiu et al., 2015).

The background activity of MNTB neurons *in vivo* are comprised of stochastic firing at ~20 Hz at steady state, plus acoustic responses that cause a higher frequency of synaptic responses, in the form of short-term plasticity, embedded in background firing (Geisler et al., 1985; Hermann et al., 2007; Liberman, 1978; Sommer et al., 1993). In our study, the EPSC steady state was also observed under the prolonged stimulation at 20 Hz with temporally Poisson-distribution (*in vivo*-like mode) or uniform 20 Hz and 50 Hz stimulation (Figures 1A–1C; Figures S1G–S1I; Figures S2A and S2B) (De Lange, De Roos, & Borst, 2003; Hermann et al., 2007). As the EPSC level at steady state corresponding to both modes of stimulation had no significant difference (Figures S2D), we determined that a uniform frequency of stimulation would be equivalent to the stochastic stimulation at the same fundamental frequency, for the purposes of studying vesicle recycling pools.

The dual whole-cell patch recording showed that prolonged 20 Hz presynaptic action potential equivalent depolarization caused a membrane capacitance increase which tended to plateau (Figure S6A). As the plateau of membrane capacitance is only reached when the exocytosis and endocytosis rates are balanced, the endocytosis could thus be estimated by the quantal content that is enhanced at PT (Figures 1F and 1G) (Renden & von Gersdorff, 2007; Smith et al., 2008). Our results confirmed the competence of the calyx of Held synapse to sufficiently supply releasable vesicles under *in vivo*-like neural activities.

Presynaptic membrane capacitance exponentially increased as much as 1.75 ± 0.15 pF from resting levels after a prolonged 20 Hz stimulation. Considering that the membrane capacitance of a single vesicle is ~ 65 fF, this amount of cumulated membrane corresponded to ~ 26,900 vesicles. This represents the first time that the surface pool of vesicle has been directly estimated, by measuring the exocytotic membrane area (capacitance) instead of assaying the fluorescence levels of a modified vesicular protein (Fernández-Alfonso et al., 2006; Hua et al., 2011; Wienisch & Klingauf, 2006). Such a large surface pool of vesicles could be sufficient to form a significant size of the readily retrievable pool (Hua et al., 2011). The existence of the surface pool also makes it possible to produce endocytic vesicles that are larger in size than exocytotic vesicles (Watanabe et al., 2014). Furthermore, the surface pool displayed obvious activity dependence, as its size varied according to the stimulation rate, strongly suggesting it functions as a regulatory stage along the recycling pathway (Figures 4A–4D). Importantly, the morphology of the ‘enlarged’ terminal under *in vivo*-like conditions, in comparison to resting conditions, should be intrinsically normal and physiological, which gives rise to the question of whether the kinetics of synaptic transmission variability corresponds to different nerve terminal sizes. To answer this question, we made presynaptic and postsynaptic dual patches and observed the kinetics of single EPSC at different time points during a train of presynaptic action potential-like stimulation. The scaled EPSCs at different times during the presynaptic membrane capacitance increase showed the same kinetics (Figures S6B–S6F). Evidently, synaptic transmission can always keep its normal characteristics, even at resting status when the terminal has ‘shrunk’. Our results show the essential robustness of synapse in transmission despite activity dependent changes in structural integrity (Kavalali & Chanaday, 2017; Xue et al., 2008).

### Kinetic dissection of recycling vesicle pool

Around 320,000 vesicles were estimated to be present in calyceal terminals by an EM study (Qiu et al., 2015). We estimated about 290,000 vesicles in the RP, ~ 90% of total vesicle in terminals, are involved in recycling. We found a great heterogeneity in kinetics of recycling among all of the RP vesicles as it took 6 ~ 10 s for a newly formed endocytic vesicle to get reused when more than 90% of the recycling vesicle remained unused (Figure S3). A key question is how the recycling SVs in the terminal are kinetically organized for synaptic transmission. A large amount of work has been done on vesicle exocytosis and endocytosis. The RRP at active zones and readily retrievable pool at the surface of nerve terminals have been proposed and well defined. Comparatively, very few studies have been done on post-endocytic vesicle trafficking (Kavalali & Chanaday, 2017; Newton, Kirchhausen, & Murthy, 2006). We previously developed a quantitative approach with exquisite signal and temporal resolution at the axo-somatic synapse and dissected the kinetics of RP depletion (Qiu et al., 2015). In this work, we studied the kinetics of vesicle trafficking along the complete recycling pathway under *in vivo*-like stimulation at physiological temperature. The RRP was estimated by back-extrapolation of cumulative EPSCs induced by a high frequency stimulation train and the surface pool was directly assayed by capacitance measurement (Figures 1H–1K; Figures 4A-4D) (Lindau & Neher, 1988; J. Y. Sun et al., 2004; Thanawala & Regehr, 2013). We found the kinetics of intra-terminal vesicle trafficking could be essentially fit by a double exponential function and well described by the transition between two sequentially connected pools, defined as the readily priming pool and the post-endocytic pool (Figures 5A and 5B) (Qiu et al., 2015). Together with exocytosis and endocytosis processes, we proposed a feed-forward four-pool model to dissect the kinetics of vesicle recycling along the whole loop of the pathway. Several justifications were made in building this model. First, we calculated and simplified the priming and de-priming process as a single feed-forward step kinetics under steady state (See Eq.22–27 in STAR Methods). Second, we derived and obtained the analytical solution from the full-parameter model based on differential equations (see Eq. 31–33 in STAR Methods). By fitting the vesicle release rate with the analytic solution, we found that backward transition from RPP to PEP should be negligible (Figures S4E–S4G). Third, we also tested a three-plus-pool model, with a hypothetic vesicle pool (PEPx) in transition to PEP, to fit the experimental results. We found that the three-plus-pool model did not significantly improve the fitness (Figures S4A and S4B). Moreover, the PEPx has much smaller size and a ten-fold faster transition rate to PEP, strongly suggesting this pool is negligible and can be merged into PEP (Figures S4C and S4D).

We found that switching the stimulation frequency from 20 Hz to 50 Hz significantly increased [Ca^2+^]_i_ and accelerated the fusion rate (k_0_), endocytic rate (k_3_) and priming rate (k_1_) (Figure 5; Figure S5) in agreement with the notion that Ca^2+^ regulates vesicle fusion, priming and endocytosis (Lou, Scheuss, & Schneggenburger, 2005; Neher & Zucker, 1993; Sakaba, 2008; Sankaranarayanan & Ryan, 2001; J. Sun et al., 2007; W. Wu, Xu, Wu, & Wu, 2005). However, the transition rate from PEP to RPP remained almost unchanged, in apparent activity independence. This is indicative and suggests the vesicles supply from PEP to RPP majorly depends on the PEP size. At this stage, the only way to meet the requirement of increasing vesicle supply in response to higher synaptic activities is to enlarge the PEP size. It is yet unknown which morphological stage corresponds to PEP and what mechanism determines this ‘bottle neck’ effect. Considering PEP is kinetically the first stage after endocytosis and a lot of endosomes and cisternae as post-endocytic structures were observed under high intensity stimulation in nerve terminals, we hypothesized that PEP could be the endocytic endosome and PEP-RPP transition possibly corresponds to the intrinsically speed limited vesicle budding and protein sorting steps from endosomes (Denzer, Kleijmeer, Heijnen, Stoorvogel, & Geuze, 2000; Hoopmann et al., 2010; McDermott & Kim, 2015; Richards, Guatimosim, & Betz, 2000; L.-G. Wu et al., 2014). It will be of great significance to associate morphological stages with these kinetic pools along the vesicle recycling pathway and thus provide more insight into the mechanism of vesicle recycling in the future.

### Kinetic structure of vesicle recycling ensures homeostasis of synaptic transmission

SVs are recycled locally to ensure neurotransmission as well as structural integrity at synapses (Denker & Rizzoli, 2010; Rizzoli & Betz, 2005; Soykan et al., 2016; Südhof, 2004). The multiple stages of vesicle recycling revealed by our study provide a wide-ranged and flexible kinetic regulation of SV recycling to support the fidelity of synaptic transmission. The nervous system is dynamic and neuronal signal is often embedded in stochastic background firing. One of the major forms of synaptic information processing is short-term plasticity which encodes transmission as a time-variant signal within milliseconds to many seconds. On the other hand, the homeostasis of synaptic transmission is required to keep the smoothness and readiness in processing instant signals with little interference by previous activity beyond a short term ‘history’. This postulation was tested by our recording of the postsynaptic EPSCs in response to stimulation with flickered frequency between 20 Hz and 50 Hz at different durations. When the postsynaptic response reached a steady state at 20 Hz, we switched the stimulation frequency to 50 Hz for different periods of time and observed the corresponding fluctuation in release rate (Figures 6A–6C). No significant difference could be found when briefly switching to 50 Hz for a duration less than 5 s (Figures 6D, 6G and 6H). Only when the switching duration was prolonged over 10 s, does the reduction in the release rate became detectable (Figures 6D–6H). It seems that the smoothness could be kept within the time window for a newly endocytic vesicle to get reused at 50 Hz stimulation (Figure S3). This homeostatic assurance could come from the kinetic structure of vesicle recycling. In our model, each feed-forward transition could be considered as a rate limiting process, mathematically represented as a homogeneous differential equation and functioning as a low pass-filter. The whole vesicle recycling pathway could be considered as an assembled series of low-pass filters which even have low cut-off frequency and filter the recent exocytotic history from endocytosis, as shown in Figure 6H. We concluded that the SV recycling machinery with the kinetic structure we have defined, could function not only as an efficient SV provider but also as a stabilizer of synaptic transmission.

## Materials and Methods

### Slice preparation and electrophysiology

Transverse slices of 200 μm thickness from 7 to 9 day old (for dual patch) or 13 to 15 day old (for long-term stimulus, presynaptic patch and calcium imaging) C57/BL6 mice were prepared with a vibratome (Leica VT 1200s). All electrophysiological experiments were performed at physiological temperature (34 °C – 36 °C) with artificial cerebrospinal fluid (ACSF) containing the following (in millimolar): 125 NaCl, 2.5 KCl, 25 NaHCO_3_, 3 myoinostol, 2 Na-pyruvate, 1.25 NaH_2_PO_4_, 0.4 ascorbic acid, 25 D-glucose, 1 MgCl_2_, and 2 CaCl_2_ at a pH of 7.4 when oxygenated (95% O_2_ and 5% CO_2_). The presynaptic pipette (3 – 4 MΩ) solution contained (in millimolar) 125 Cs-gluconate, 20 CsCl, 4 MgATP, 10 Na_2_-phosphocreatine, 0.3 GTP, 10 Hepes, and 0.05 BAPTA (pH 7.2 adjusted with CsOH). The postsynaptic pipette (2 – 3 MΩ) solution contained (in millimolar) 125 K-gluconate, 20 KCl, 4 MgATP, 10 Na_2_-phosphocreatine, 0.3 GTP, 10 Hepes, and 0.5 EGTA (pH 7.2 adjusted with KOH). Presynaptic and postsynaptic whole-cell recording were obtained with an EPC-10 amplifier (HEKA) and an Axopatch 200B amplifier (Axon Instruments), with series resistances < 15 MΩ and 10 MΩ compensated by 60% and 95% (lag 10 μs), respectively. The holding potential of − 80 mV was corrected for a liquid junction of − 11 mV between the extracellular and pipette solution. For dual-patch experiments, tetrodotoxin (TTX; 1 μM) and tetraethylammonium (TEA; 20 mM, replacing 20 mM NaCl) were added to block Na^+^ and K^+^ channels, and cyclothiazide (CTZ; 0.1 mM) were applied to prevent postsynaptic EPSC saturation. Afferent stimuli were delivered by electrical stimulation (0.1 ms, 3 – 30 V) via a bipolar electrode positioned at the midline of the trapezoid body and EPSCs were recorded in the presence of CTZ (0.1 mM, all experiments). Folimycin (Calbiochem) was first dissolved in DMSO and then in ACSF. Stock solution (57.7 μM) was prepared and stored at − 80 °C. The drug was diluted in ACSF to a final concentration of 4 μM on the day of use. The final DMSO concentration was < 0.3%. The drug was applied to the patched cells by puffing at a distance of less than one cell diameter via a Picospritzer III microinjector. Data are expressed as the mean ± SEM and student’s t-test were used. All data were analyzed with Igor Pro-6.2 (WaveMetrics). The care and use of mice in all experiments conformed to institutional policies and guidelines (IBP animal protocol 8701, approved by the Ethics Committee of Chinese Academy of Sciences Institute of Biophysics).

### Ca^2+^ imaging

Experiments involving Ca^2+^ detection were made by loading Fura-4f (200 μM). The cells were then excited at 340 nm using a polychromiatic illumination system (T.I.L.L. Photonics, Munich, Germany). Images were collected and off-line analyzed by Andor iXonEM ^+^ hardware and software system. Changes in cytosolic Ca^2+^ ([Ca^2+^]_i_) for Fura-4f loaded cells were measured by converting the ΔF/F_0_ ratio of fluorescence (after background subtraction) to approximate [Ca^2+^]_i_, corresponding to 20 Hz/50 Hz action potential-like stimulations.

### Estimation of mEPSC amplitude in the presence of folimycin

Based on the fact that folimycin reduced the vesicular transmitter content without affecting the fusion machinery or the vesicle release probability, the fractional change in evoked EPSCs would reflect the fractional change in quantal size when newly endocytosis vesicles were fully reused. Thus, the size of mEPSCs with folimycin (q_foli_) is estimated as

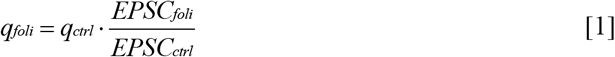

Where q_ctrl_ is the average mEPSC amplitude during the resting period before folimycin application. EPSC_foli_ and EPSC_ctrl_ represent the average amplitude of the last 100 EPSCs in the folimycin and control groups, respectively.

### Kinetics of vesicle recycling

We attributed the cumulative EPSC amplitude to the summative preserved SVs (N_preserved_) with qctrl and reused SVs (N_reused_) with q_ctrl_ (control group) or q_foli_ (folimycin group) (Qiu et al., 2015):

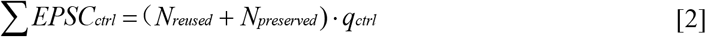

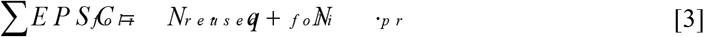

Where ∑ EPSC_ctrl_ and ∑ EPSC_foli_ represent the cumulative EPSC amplitudes in the control and folimycin groups, respectively. Thus, N_preserved_ and N_reused_ can be derived as:

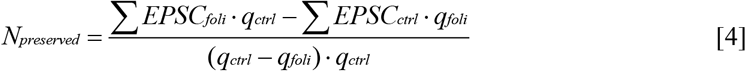

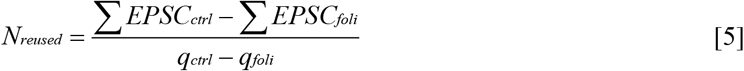

### Three-pool kinetic model

In the model, we have the general assumption that vesicle release during the steady state phase is limited by the constant rate of vesicle supply, the RP depletion rate (R_depl_), assayed as release rate, is approximately the same as the priming rate as described in Eq. 6–7. By designating RPP(t) and FP(t) as the pool sizes at a given time t, the relation among R_depl_, RPP, and FP are represented as:

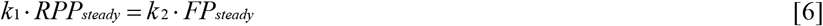

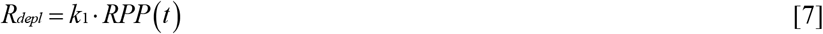

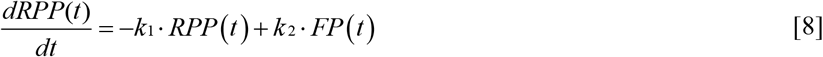

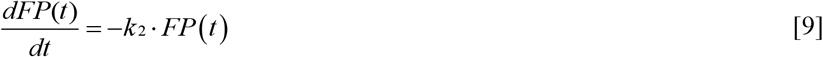

Where k_1_ is the formal priming rate and k_2_ is the supply rate from FP to RPP.

The analytic solution was derived by solving the following transformed differential equation:

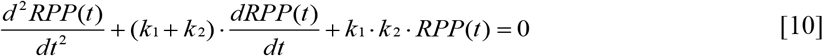

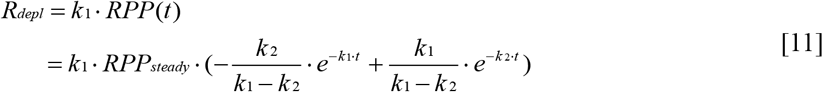

Where 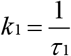 and 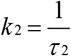.

In case k_1_ = k_2_ ≈ k the analytic solution could be represented as:

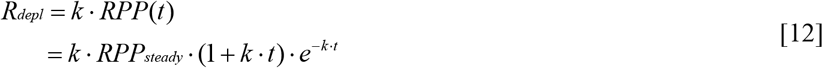

Where 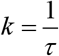

All the kinetic parameters were estimated by fitting the measured *R_depl_* with a double-exponential function or mono-exponential function (the case with two very close time constants) as the analytic solution.

### Three-plus-pool kinetic model

In assumption that two sequentially connected pools (FP1 and FP2) followed the RPP, we proposed a three-plus-pool kinetic model, represented as:

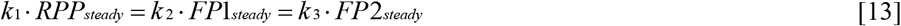

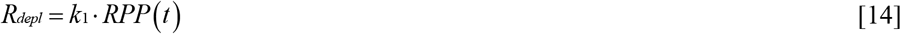

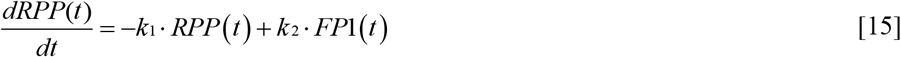

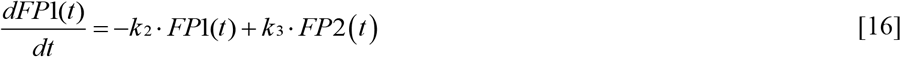

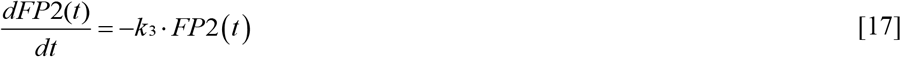

Above equations could be derived as

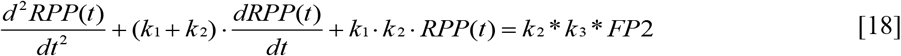

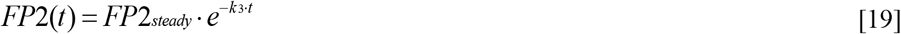

For k_1_ ≠ k_2_, the analytic solution of is:

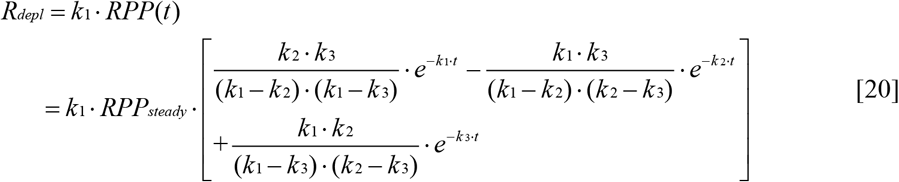

For k_1_ = k_2_ ≈ k, the analytic solution of is

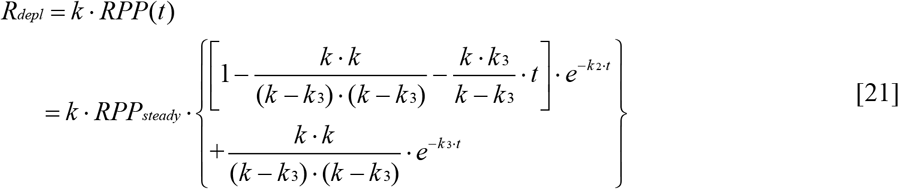

RPP_steady_ and FP_steady_ can thus be obtained by fitting the measured R_depl_ with the above analytic solutions.

### Full-parameters model including forward and backward reactions

If the forward and backward transition among each step of recycling was taken into account, the full-parameter model can be described as:

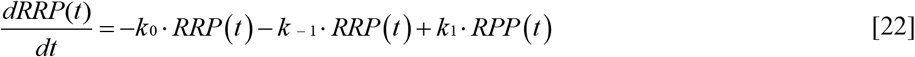

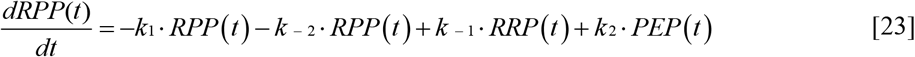

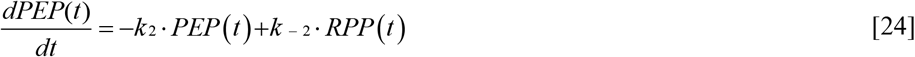

At the steady state, as generally assumed, 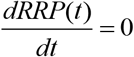

Equation [22] can be derived as

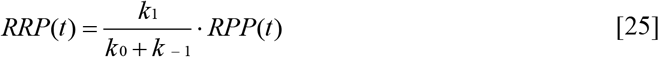

Equation [23] can be reorganized as

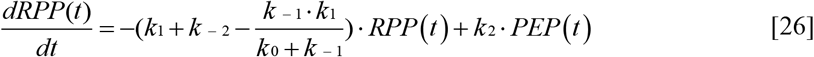

Let

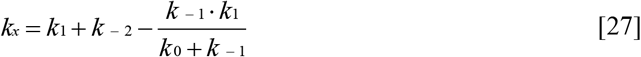

we can rewrite the last two equations as

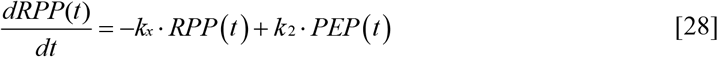

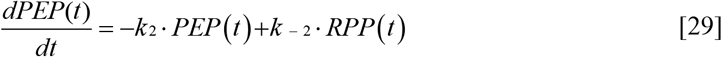

The general solution of this differential equations can be solved by:

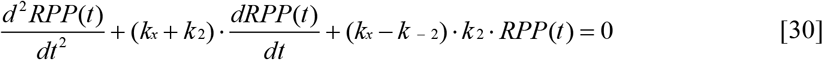

The analytical solution is expressed as

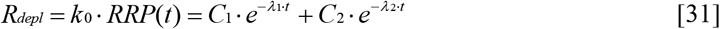

Where

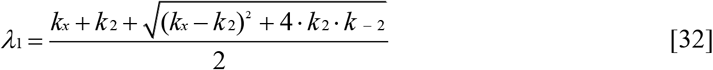

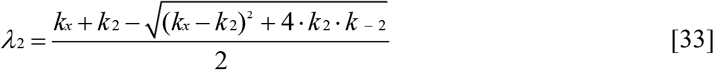

The R_depl_ is fitted by the analytic solution of these equations to obtain the parameters of k_2_ and k^−2^.

## Author contributions

Zili Liu: Conceptualization; Investigation; Writing - original draft. Ying Zhu: Conceptualization; Investigation; Writing - original draft. Yubing Hu: Investigation. Jianyuan Sun: Conceptualization and project design; Supervision; Writing – editing and revising

## Data Availability

All data generated or analyzed during this study are included in the manuscript and supporting files.

## Ethics

Human Subjects: No. Animal Subjects: Yes. Ethics Statement: The care and use of mice in all experiments conformed to institutional policies and guidelines (IBP animal protocol 8701, approved by the Ethics Committee of Chinese Academy of Sciences Institute of Biophysics).

## Acknowledgments

We thank Drs. T. Südhof, E Neher, Ken Paradiso, Z. P. Pang, J Sorensen for beneficial discussions and critical comments on the manuscript; H. Tian for support with mEPSC analysis; and X. D. Zhao and S. Liu for technical support. This work was supported by the National Natural Science Foundation of China (31871033 to J.S. and 31527802 to J.S.), the National Basic Research Program of China (2013CB835100 to J.S.), the Instrument Development Project, CAS (YJKYYQ20180028 to J.S.), and the Key Research Program of Frontier Sciences, CAS (QYZDYSSW-SMC015 to J.S.).

## Author Contributions

Z.L., Y.Z. and J.S. designed the research; Z.L. and Y.Z. performed experiments; Z.L., Y.Z. and Y.H. analyzed data; Z.L., Y.Z. and J.S. wrote the paper.

## Supplemental Information

**Figure S1.**
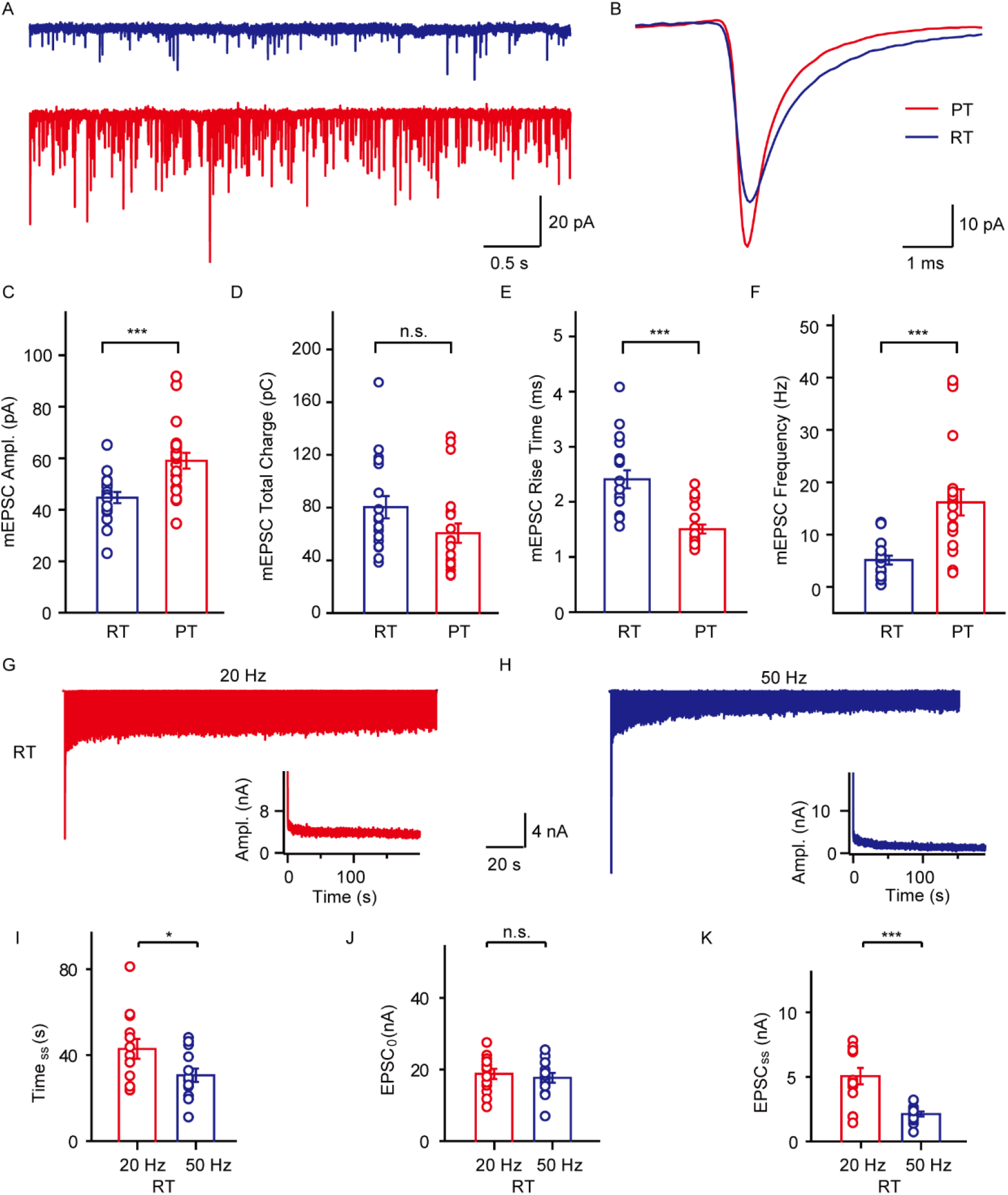
Comparison of mEPSCs at RT and PT and the steady state of synaptic transmission at RT. (A) Representative recorded mEPSC traces at RT (Blue) and PT (Red). (B) The averaged single mEPSC traces at RT (Blue) and PT (Red). (C-F) Statistics of mEPSC amplitude (C), total charge (D), rise time (E) and frequency (F) (RT: n = 18; PT: n = 21). (G and H) Representative EPSCs and the corresponding amplitudes during sustained stimulation at 20 Hz (G) or 50 Hz (H) at room temperature. (I-K) Statistics of steady state time (I), first EPSC amplitude (J), and steady state amplitude (K) of each train at 20 Hz or 50 Hz stimulation at room temperature (n=13). Data are represented as mean ± SEM. \**p* < 0.05, \*\*\**p* < 0.001.

**Figure S2.**
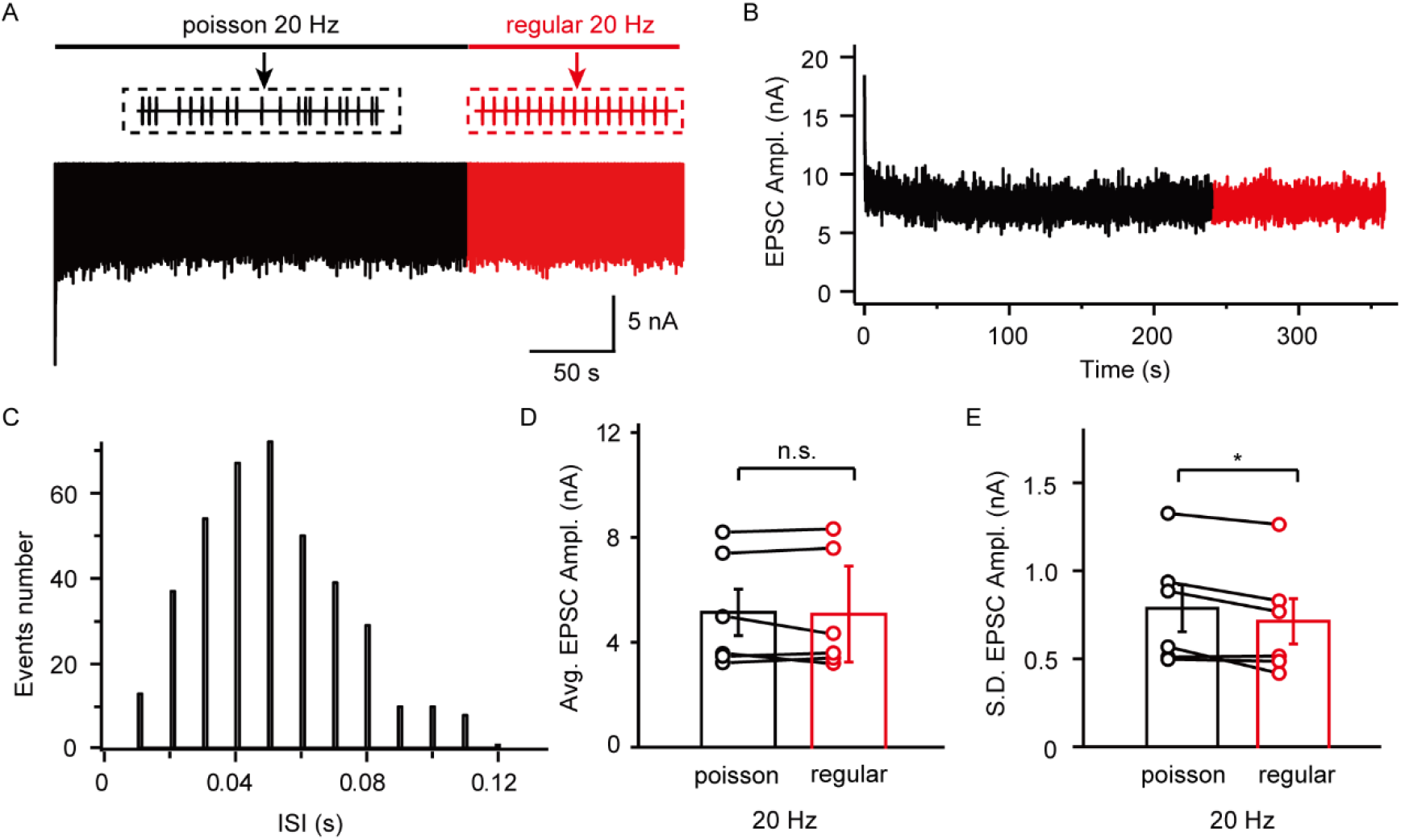
EPSCs under temporal Poisson-distribution frequency stimulation and constant frequency stimulation. (A and B) Representative EPSCs (A) and corresponding amplitude (B) evoked with the afferent fiber stimulation sequence with 240 s of the Poisson distribution and 120 s uniform stimulation at fundamental frequency of 20 Hz. (C) The time interval of Poisson distribution generated by pseudo-random sequence. (D and E) Statistics of the EPSC amplitudes (D) and amplitudes standard deviation (E) at steady state under Poisson distribution stimulation and uniform stimulation. Data are represented as mean ± SEM and n = 7. *\*p* < 0.05.

**Figure S3.**
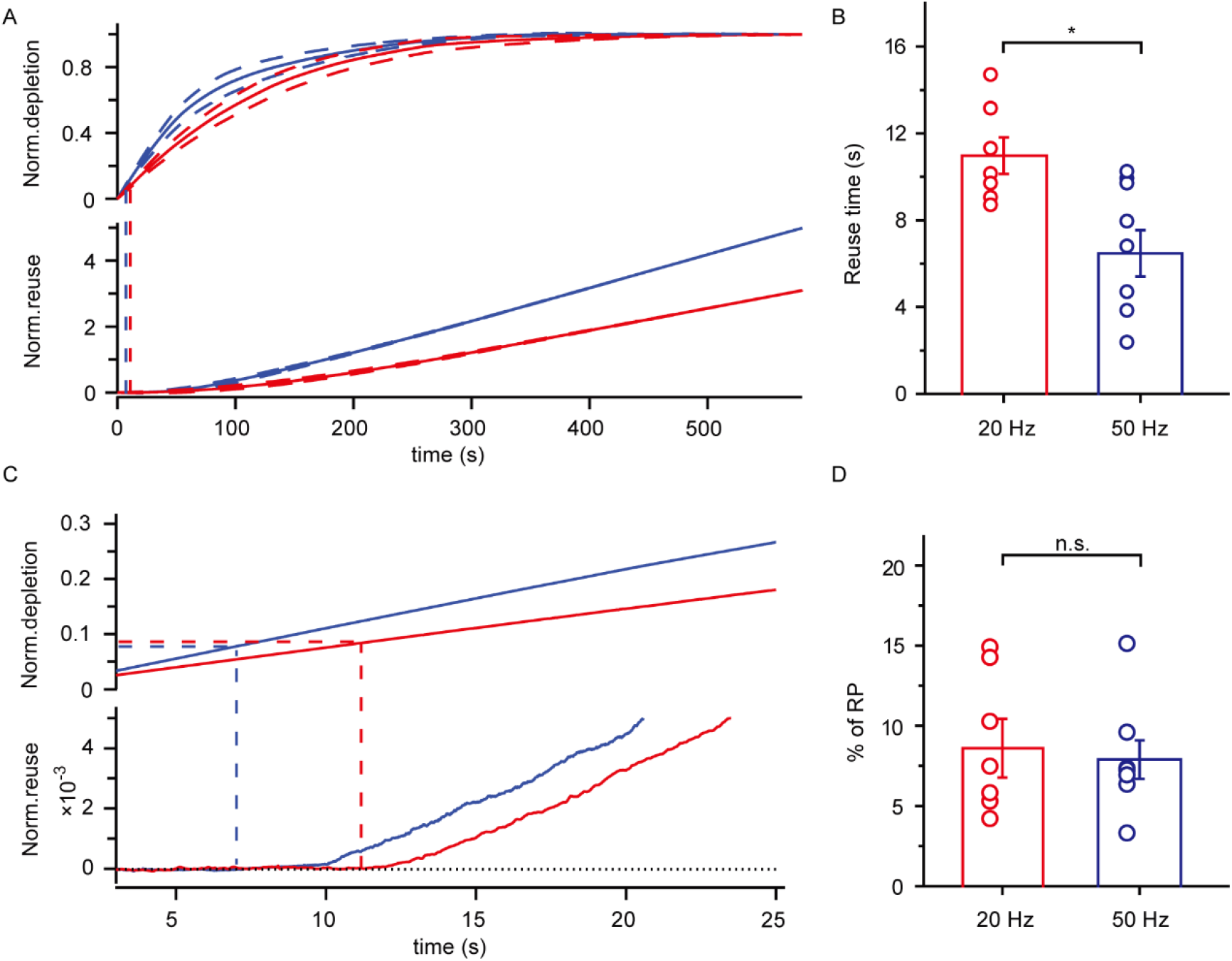
Timing of vesicle reuse under sustained stimulation. (A) Cumulative preserved SV (Upper) and reused SV (Lower) numbers normalized to preserved pool size under 20 Hz (blue, n = 7) and 50 Hz (red, n = 8) stimulations. The dashed lines denote mean ± SEM. The vertical dashed lines indicate the starting time points of vesicle reuse. (B) Statistics of starting time of reuse under 20 Hz and 50 Hz stimulation; * *p* < 0.05, t. tests. (C) The local zoom of starting time to reuse in (A) (D) Statistics of the fraction of RP depletion at the starting time point of reuse under 20 Hz and 50 Hz stimulation. Data are represented as mean ± SEM. \**p* < 0.05.

**Figure S4.**
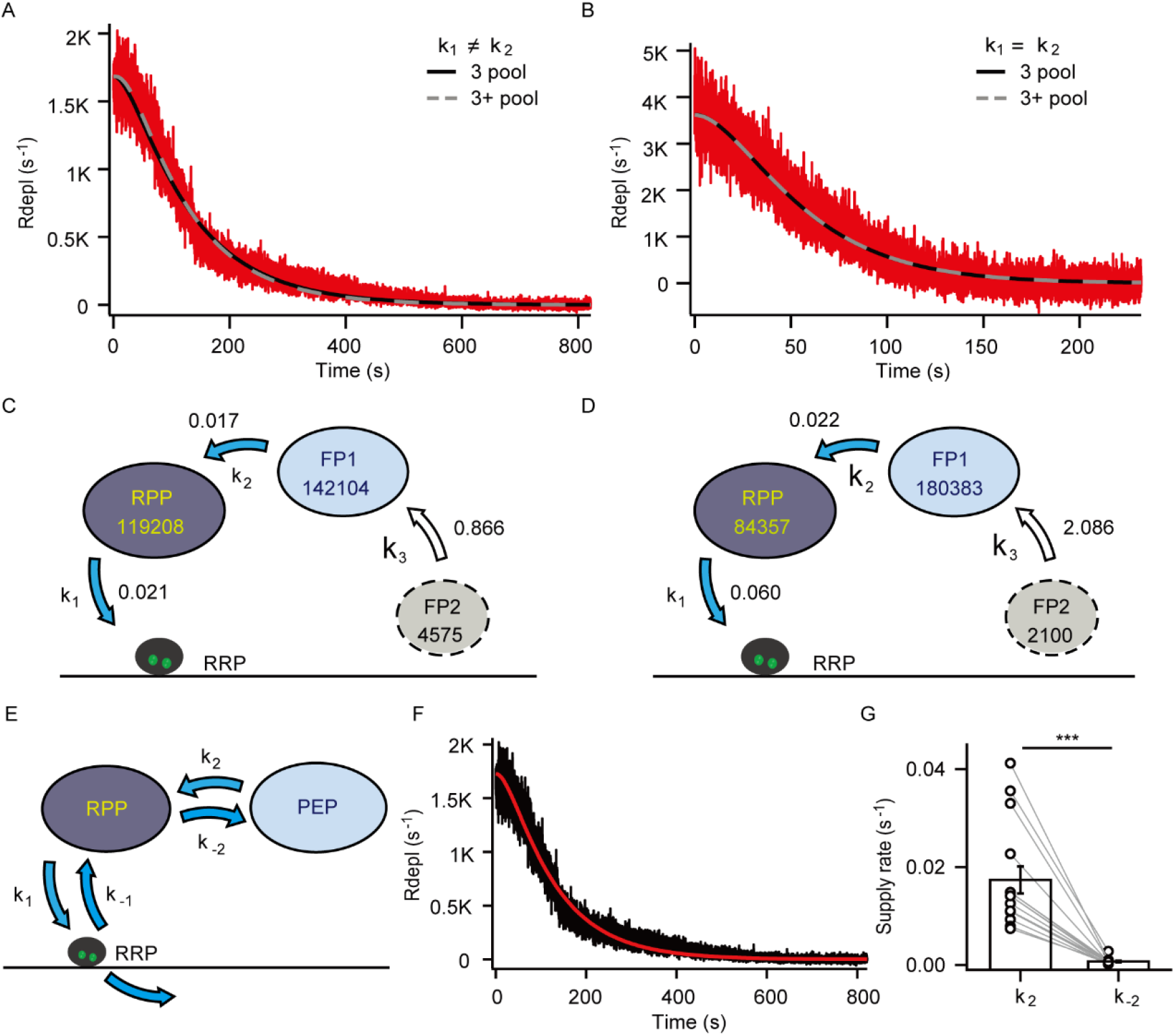
Three-plus-pool model and full-parameters model of preserved pool depletion. (A and B) Representative preserved pool depletion rate (Rdepl) trace with sequential three-plus-pool model fitting and three-pool model fitting with k_1_ ≠ k_2_ (A) and k_1_ = k_2_ ≈ k (B). (C and D) Quantification of dissected synaptic vesicle pools by three-plus-pool model under 20 Hz stimulation (n = 7) (C) and 50 Hz stimulation (n = 8) (D). (E) Schematic of full-parameters model of preserved pool depletion. (F) Representative preserved pool depletion rate (R_depl_) trace with full-parameters model fitting. (G) Parameters of k_2_ and k_-2_ were obtained by fitting the measured R_depl_. k_2_ (0.0174 ± 0.0028) is much larger than k_-2_ (0.0007 ± 0.0002) from measured cells (n = 15). Data are represented as mean ± SEM. \*\*\**p* < 0.001.

**Figure S5.**
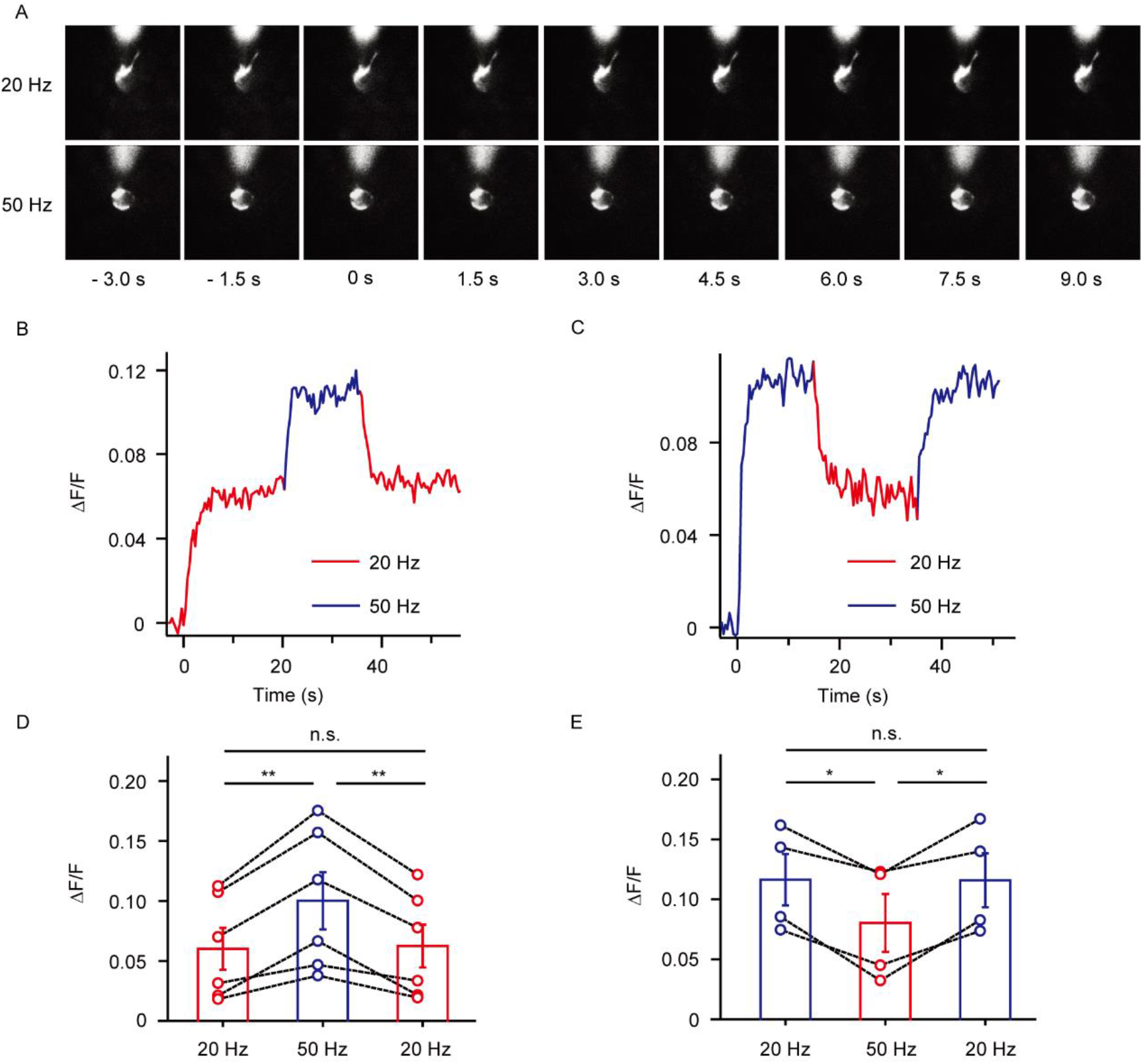
The intracellular calcium concentration under sustained stimulation. (A) The representative process of Ca^2+^ indicator Fura4f fluorescence in the synaptic terminal at calyx of Held synapse during depolarization at 20 Hz and 50 Hz from resting status. (B and D) The representative fluorescence trace of the Fura4f under 340 nm excitation light evoked by the continuous depolarization of 20 Hz-50 Hz-20 Hz (B) and the corresponding statistical results (n = 6) (D). (C and E) The representative fluorescence trace of the Fura4f under 340 nm excitation light evoked by the continuous depolarization of 50 Hz-20 Hz-50 Hz (C) and the corresponding statistical results (n = 4) (E). Data are represented as mean ± SEM. \**p* < 0.05, \*\**p* < 0.01.

**Figure S6.**
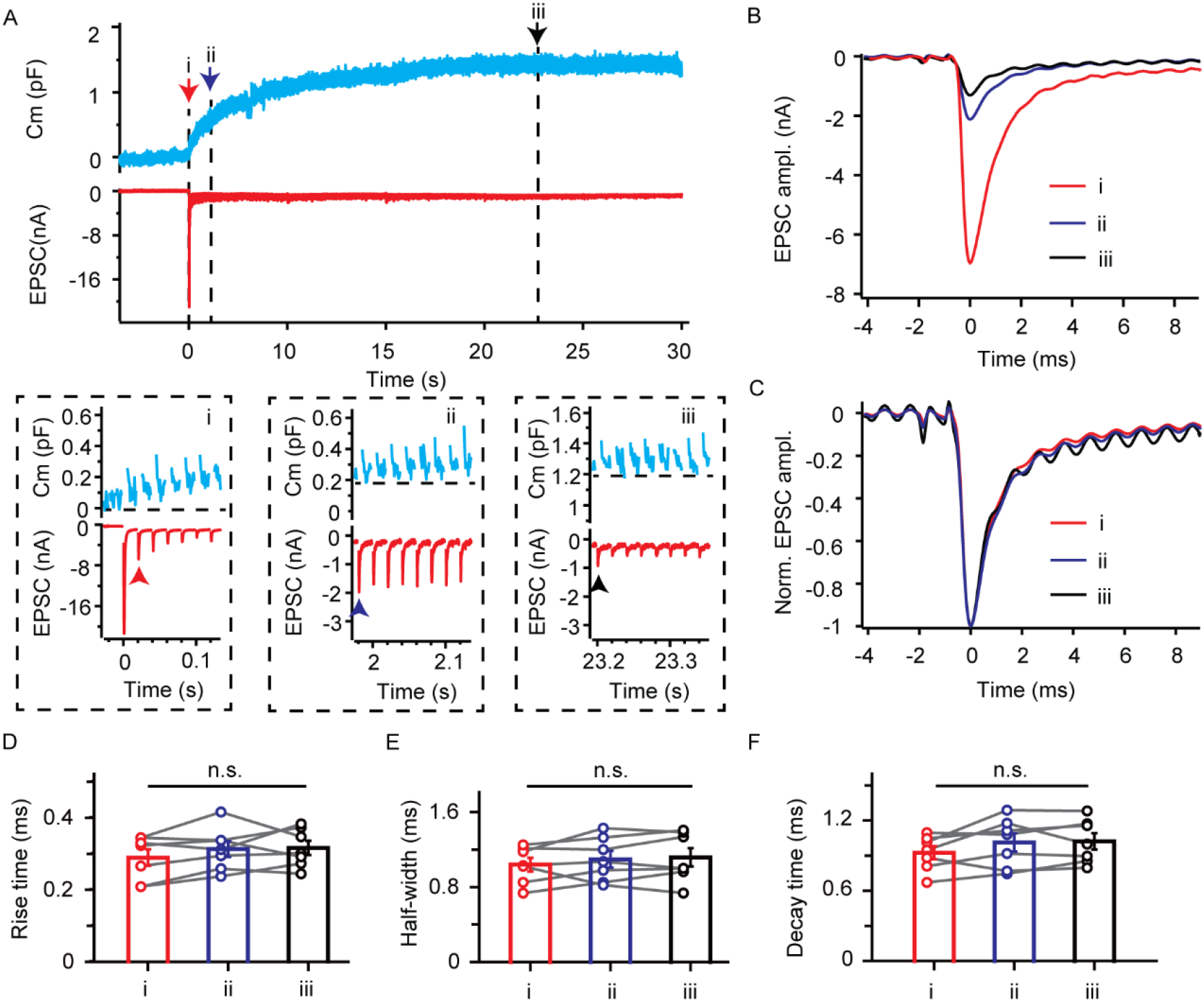
Kinetics of EPSCs with different sizes of surface pool during sustained stimulation. (A) The representative capacitance measurement and EPSCs recording during 20 Hz presynaptic AP-equivalent depolarization and local zoom at different time points of presynaptic membrane capacitance increase. (B and C) The recorded EPSCs (B) and scaled EPSCs (C) at different time points of presynaptic membrane capacitance. (D-F) Statistics of rise time (D), half-width (E) and decay time (F) at different time points of presynaptic membrane capacitance. Data are represented as mean ±SEM and n = 7.

